# D1R- and D4R-Dependent Regulation of Retinal Light Adaptation in Diabetes

**DOI:** 10.64898/2025.11.29.691314

**Authors:** David Umbertus, Angela MacIsaac, Erika D. Eggers

**Affiliations:** University of Arizona

## Abstract

Diabetic retinopathy (DR) is considered a neurovascular disease, and key clinical features show impairments in visual acuity and electroretinogram (ERG) recordings. Dopamine, a critical regulator of light adaptation, is also reduced in diabetes and supplementation of dopamine or injection of dopamine agonists improve visual deficits in diabetic rodents and humans. We sought to examine how blocking D4Rs and D1Rs would affect ERG signaling in 6-week diabetic rodents. Blocking D4Rs with L745,870 significantly reduced dark-adapted A-wave amplitude in DM, but not non-DM retinas, with no change in a-wave implicit timing, suggesting diabetic alterations in D4R signaling in photoreceptors. Light-adapted (LA) recordings revealed blocking D4Rs significantly increased A-wave amplitude and delayed A-wave implicit timing in DM and non-DM retinas. B-wave amplitudes were elevated at the highest flash intensity, and rise time was significantly faster, indicating an inability to adjust to background light. D1R blockade with SCH-23390 had little effect on ERG recordings for both groups, aside from increased B-wave amplitude and delayed rise time in dark-adapted DM conditions towards the lower light intensities. While small, D4R blockade seems to have a bigger effect in DM conditions during light adaptation, suggesting that diabetes impairs the retina’s ability to adjust to light adaptation. These studies show that dopamine signaling in diabetes is perturbed and an important avenue for future investigation.

## 1. Introduction

Diabetic retinopathy (DR) was prevalent in about 27% of individuals diagnosed with diabetes mellitus (DM) in 2021 equating to 9.6 million individuals in the US^[1-4]^. DR is classically defined as a microvascular disease with two stages: non-proliferative DR (NPDR) and proliferative DR (PDR). Patients with DR have reported impairments in visual acuity and contrast sensitivity, disrupted color vision, and slowed dark adaptation^[5-8]^. However, prior to DR vascular damage, electroretinogram (ERG) recordings show deficits in neuronal signaling as early as 4 weeks after the onset of diabetes in rodents^[9, 10]^. Furthermore, multifocal ERGs have detected different sites of diminished electrical activity across the retina predicting future sites of vascular development^[3, 11, 12]^.

Another feature of diabetes is reduction of retinal dopamine (DA)^[13-15]^ as early as 5-weeks after diabetic onset^[14]^ in mice. Supplementing diabetic mice and rats with dopamine and dopamine agonist injections improved visual deficits and ERG deficits^[14, 15]^. Similar outcomes were found in diabetic humans^[16]^, and emphasize the importance of understanding dopamine signaling dysfunction in the diabetic retina.

Dopamine is a critical regulator of the retina’s ability to adjust a wide range of light intensities, otherwise known as light adaptation (LA). This process is important for normal visual function and is regulated by dopamine release from dopaminergic amacrine cells^[17-19]^. Dopamine binds to dopamine receptors (R) on photoreceptors (D4Rs), horizontal/bipolar/amacrine/ganglion cells (D1Rs), and dopaminergic amacrine cells (D2Rs)^[20-28]^. Activating D1Rs and D4Rs reduces ganglion cell responses^[29]^ and after 6-weeks of diabetes, ON-sustained ganglion cells had a reduced ability to adjust to increasing levels of background light^[30]^. This is possibly attributable to reductions in dopamine D4R sensitivity^[31]^ and likely not receptor expression as mRNA expression of D4R and D1Rs did not change in the diabetic mouse^[13, 14]^.

Light adaptation relies on dopamine to facilitate photoreceptors and downstream circuitry to adequately adjust their sensitivity to increasing background luminance. In photoreceptors, D4R activation reduces the amount of cAMP through adenylyl cyclase (AC) suppression, while D1R activation increase the amount of cAMP and modulates the gain control of horizontal, bipolar, amacrine, and ganglion cells. Therefore, we would assume that applying D4R/D1R antagonists would disrupt the adaptation process. We have recently shown that application of either D4R or D1R agonists had no effect on ERG responses in 6-week diabetic mice^[13]^ suggesting that there may be underlying changes in dopamine receptor activity or protein levels in diabetes. Because of this, the aim of our study was to identify ERG abnormalities by blocking D4Rs and D1Rs in early diabetes and to determine if blocking these receptors limited light adaptation or if diabetes had limited responsiveness to dopamine receptor antagonists. We analyzed dopamine signaling changes using ex vivo whole retina ERG recordings and compared non-diabetes mellitus (non-DM) versus diabetes mellitus (DM) groups 6 weeks after injections.

## 2. Methods

### 2.1 Animals

All animal protocols were approved by the University of Arizona’s Institutional Animal Care and Use Committee and followed the ARVO Statement for the Use of Animals in Ophthalmic and Vision Research. 5-week-old C57BL/6J male mice (Jackson Laboratories, Bar Harbor, ME) were fasted for 4 hours and then injected intraperitoneally with either streptozotocin (STZ; 75mg/kg body weight) in 0.01M sodium citrate buffer (pH 4.5) for the diabetic group (DM), or citrate buffer for the non-diabetic group (non-DM). Blood glucose was measured (One Touch Ultra Mini; LifeScan, Milpitas, CA) 6 weeks after injections in mice fasted for 4 hours (Table 1). STZ-injected mice with blood glucose <250 mg/dl and citrate-injected mice with blood glucose >250mg/dl were excluded from the study. Mice were given access to food (NIH-31 rodent diet) and water ad libitum.

### 2.2. ERG Procedures

#### 2.2.1. Drugs and Solutions

A modified Ringer’s solution (125mM NaCl, 2.5 mM KCl, 1 mM MgCl2, 1.25mM NH2PO4, 20mM Glucose, 26mM NaHCO2, 2mM CaCl2, 2 mM Na-Pyruvate, 4mM Na L-Lactate, 0.5 mM L-Glutamine), bubbled with 5% CO2-95% O2 to maintain pH ∼7.4, was used to mimic extracellular fluid. Barium chloride (100μM) was added the morning of each experiment to block electrical potential from Müller cells (Green and Kapousta-Bruneau, 1999). D4R antagonist L745,870 trihydrochloride (L745, 1 μM, Tocris, Bristol, UK) and D1R antagonist SCH-23390 hydrochloride (SCH, 50 μM) were used to inactivate D4Rs and D1Rs, respectively. All drugs were purchased from Sigma-Aldrich (St. Louis, MO) unless noted otherwise.

#### 2.2.2. Retina Preparation and ERG Setup

All ERG Dissections, preparations, and recordings were performed under infrared illumination to preserve light sensitivity of the retinal tissue. 6 weeks following STZ or citrate injections, mice were dark-adapted overnight, then euthanized with carbon dioxide. Eyes were enucleated and placed into bubbled cold extracellular solution (See section 2.2.1). The cornea and lens were removed, the retina was dissected from the eyecup, and the retina was transferred to 0.4 μm PCF filter paper (Millipore, Billerica, MA). The retina was mounted photoreceptor side up and filter paper was placed into a custom 3D printed ERG chamber (PTFE filament) which was then filled with cold extracellular solution.

The chamber was connected to an *ex vivo* ERG and extracellular solution was bubbled and perfused into the chamber to ensure the tissue remained oxygenated (VCS Computer-controlled Valve controlled System; Smart Ephys, Harvard Bioscience). The bath level was maintained by modifying the outflow (Fisherbrand Variable-Flow Peristaltic Pump; Thermo Fisher Scientific, Waltham, MA). The bath temperature was maintained within a range of 35° C to 37° C (TC-324B temperature controller with SHM-6 multi-line in-line solution heater; Warner Instruments, Holliston, MA).

#### 2.2.3. ERG Recordings

Full-field light stimuli were produced by a light emitting diode (LED; NTE30066, λpeak = 523 nm; NTE Electronics Inc) calibrated with an S471 optometer (Gamma Scientific, San Diego, CA). Light was projected through the camera port of an Eclipse FN1 Microscope (Nikon Instruments, Tokyo, Japan) onto the chamber through a 4x objective. Signals were measured using Ag-AgCl pellet electrodes (E210, Warner Instruments). Recordings were sampled at 10kHz, filtered with a 1kHz low-pass filter (DP-311 Differential Amplifier; Warner Instruments) and digitized with Axon Digidata 1550B (Molecular Devices, San Jose, CA). Clampex 11.2 software (Molecular Devices) controlled stimulus intensity (3.88s105 photons.μm™2.s™1), duration of light stimulus (30ms), duration of intervals (20s) and number of sweeps per recording set (15). The retina was equilibrated while being perfused with extracellular solution in the ERG setup for 15 min before stimuli were applied We ran two dark-adapted (DA) recording sets with intensity flashes at 9.5 photons/µm^2^/s, 950 photons/µm^2^/s, 9.5*10^3^ photons/µm^2^/s, 9.5*10^4^ photons/µm^2^/s, and 9.5*10^5^ photons/µm^2^/s twice before either the D1R or D4R antagonist solution was added. D1R or D4R antagonist was then perfused for 15 min and two DA light-stimulated recording sets were repeated for both non-DM and DM groups. Following this, we performed our light-adapted recording set that introduced a background light intensity of 950 photons/µm^2^/s with three light stimuli at three increasing intensities: 9.5*10^3^ photons/µm^2^/s, 9.5*10^4^ photons/µm^2^/s, and 9.5*10^5^ photons/µm^2^/s.

#### 2.2.4. ERG Data Analysis and Statistics

Recordings were analyzed using Clampfit software (Molecular Devices) to extract the peak amplitudes and implicit times for each peak and are described in MacIsaac et al, 2024. Implicit time to peak was calculated from the time of the light stimulus to the minimum or maximum value of a peak. A-wave amplitude was calculated from the baseline mean value to the minimum value of the A-wave. B-wave amplitude was calculated from the minimum value of the A-wave to the maximum value of the B-wave. Amplitudes were normalized to the maximum value in control solution for individual retinas to account for the differences in responsivity between retinas. Rise time was calculated from the time of minimum value in the A-wave to the time of the maximum value in the B-wave. Data from individual stimulus responses was averaged for each retina and either 2-way ANOVA or mixed-effects ANOVAs were run using the Tukey method of pairwise comparison using GraphPad (Insight Partners, NYC, New York). Plots were graphed using the GraphPad (Insight Partners, NYC, New York). All differences were considered significant if P < 0.05.

## 3. Results

### 3.1 Dark-adapted A-wave amplitudes are reduced in DM retinas after application of D4R antagonist

We first investigated how blocking dopamine receptors (D4R/D1R) affected ERG waveforms after 6 weeks of diabetes. We used L745,870 trihydrochloride (1 μM; L745) to antagonize D4Rs that are primarily expressed on photoreceptors. Light responses were recorded from dark-adapted retinas before and after application of the D4R antagonist, L745, in non-DM and DM groups. We normalized retinas to their maximum intensity response without drug application to isolate drug effects by removing variability in retinal responsivity. In non-DM retinas, we observe a slight decrease, though not significant, in A-wave amplitude after application of L745 (2-way ANOVA p = 0.16, Non-DM C vs. L745 Tukey, Figure 1C). However, we do see a significant decrease in A-wave amplitude in DM groups after application of L745 (2-way ANOVA p=0.0273; DM C vs L745 Tukey, Figure 1C), and this is reflected in the example trace (Figure 1B) where the a-wave peak in the DM-L745 group (orange) is smaller than that of the DM group (black).

**Figure 1.**
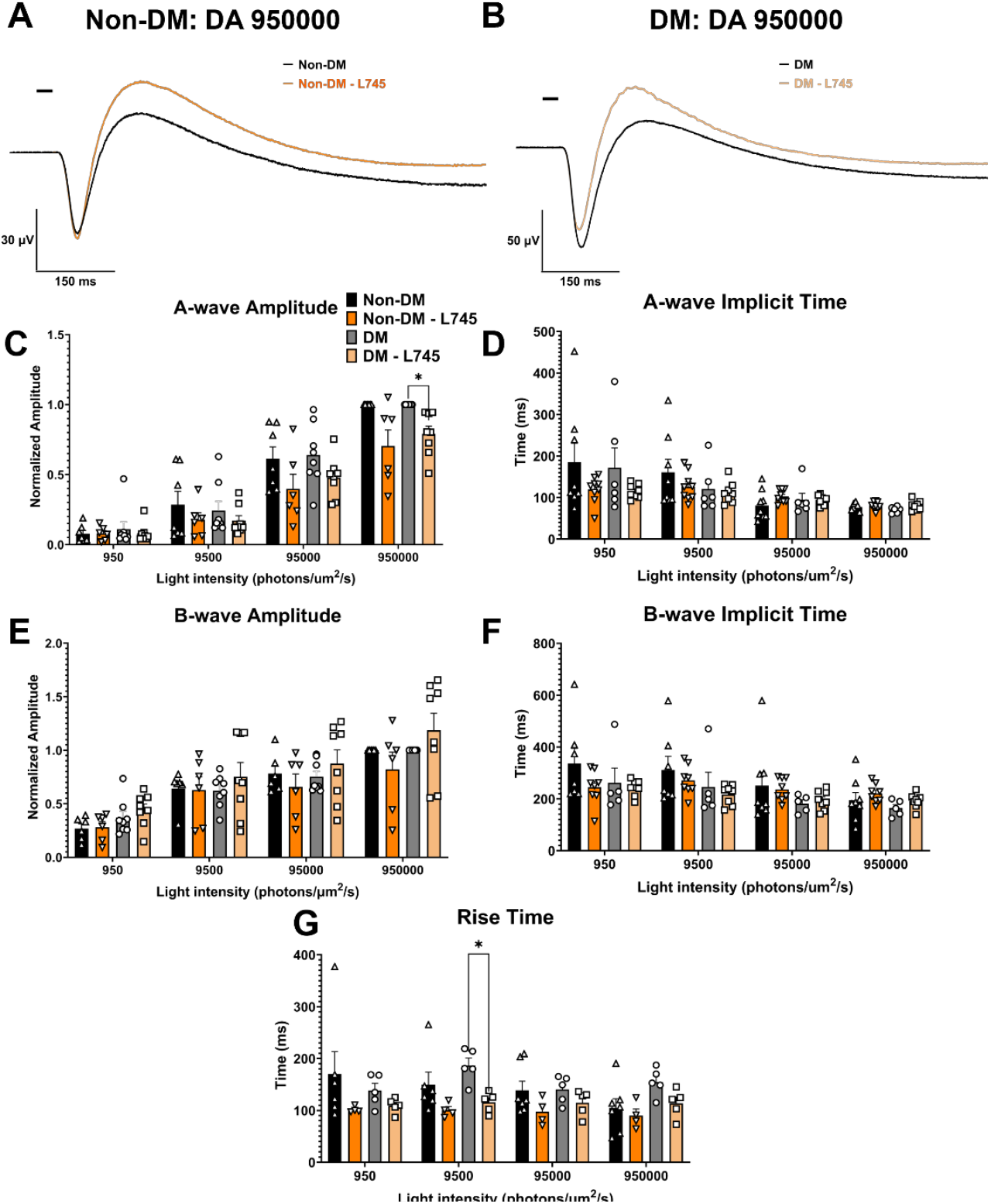
D4 receptor antagonism selectively reduces dark-adapted A-wave amplitude in diabetic retinas. (A-B) Representative dark-adapted ERG traces from non-diabetic (Non-DM) and diabetic (DM) retinas before (black) and after L745,870 application (1 μM; non-DM, orange; DM, light orange). (C) Quantification of A-wave amplitude shows no significant change in Non-DM retinas but a significant reduction in DM retinas after L745. (D) A-wave implicit time was unchanged in both groups. (E-F) B-wave amplitude and implicit time were not significantly affected by L745 in either condition. (G) Rise time was significantly faster in DM retinas at the highest flash intensity following L745 application. Data are shown as mean ± SEM with individual retinas plotted. Statistical comparisons performed using 2-way ANOVA with Tukey post hoc tests.

These decreases had no significant effect on implicit timing from the onset of the flash to the peak of the A-wave (Figure 1D). For both groups, there are no significant changes related to the amplitude (Figure 1E) or implicit timing to the b-wave (Figure 1F). We do see a significant decrease in rise time at the flash intensity of 9500 photons/um^2^/s in the DM group after application of L745 (2-way ANOVA p = 0.0178, DA 9500 DM C vs. L745 Tukey, Figure 1G). These results suggest that blocking D4Rs alters A-waves in DM retinas versus non-DM retinas.

### 3.2 Light-adapted ERG responses are altered after application of D4R antagonist

We then explored LA and how this differed between non-DM and DM groups with application of L745. In normal LA (without drugs), A-wave and B-wave amplitudes are reduced to roughly 50% of their maximal dark-adapted responses (Figure 2C, E; 950,000 photons/um^2^/s), reflecting their ability to reduce their sensitivity. With application of L745, both non-DM and DM groups had significant increases in the A-wave amplitude (2-way ANOVA, p = 0.0010, non-DM C vs. L745 Tukey; 2-way ANOVA p = 0.0048, DM C vs. L745 Tukey, Figure 2C) and we consistent delays in a-wave timing in both conditions (Mixed-effects analysis p = 0.0205 (9500); 0.0005 (95000);

**Figure 2.**
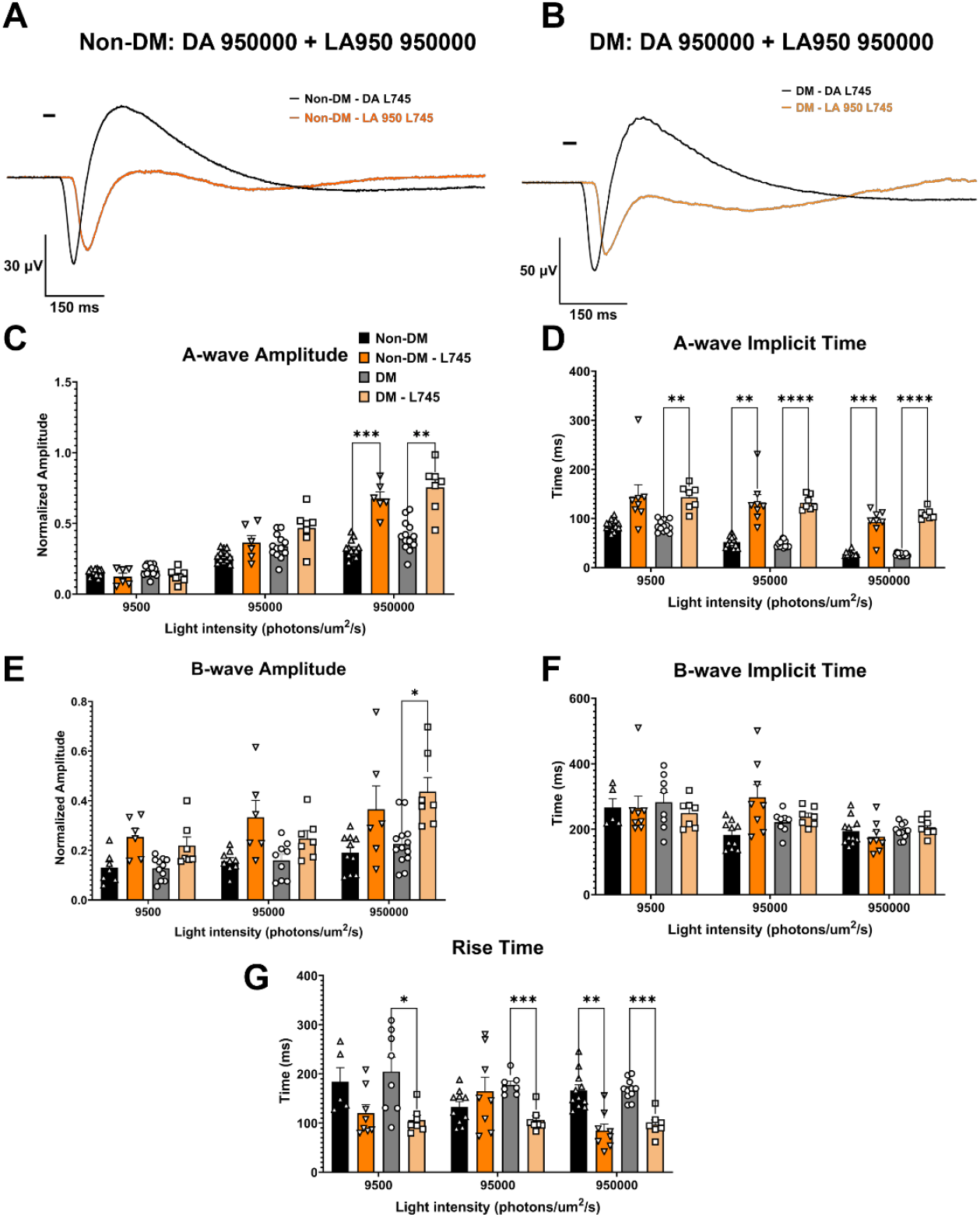
D4 receptor blockade disrupts light adaptation and delays A-wave timing in both Non-DM and DM retinas. (A-B) Representative light-adapted ERG traces for Non-DM and DM retinas before (black) and after L745 application (non-DM; orange and DM; light orange). (C) A-wave amplitude significantly increased with L745 in both Non-DM and DM groups at highest flash intensity. (D) A-wave implicit time was significantly delayed following D4R blockade, with exaggerated delays in DM retinas. (E) B-wave amplitude was largely unaffected except for a significant increase at the highest flash intensity in both groups. (F) B-wave implicit time did not change with L745. (G) Rise time was significantly faster after L745 treatment in DM retinas at all intensities, and in Non-DM retinas at the each intensity, indicating impaired adaptation kinetics. Mean ± SEM with individual data points; mixed-effects or 2-way ANOVA with Tukey post hoc tests as appropriate.

0.0007 (950000), non-DM C vs. L745 Tukey; Mixed-effects analysis, p = 0.0197 (9500); 0.0003 (95000); 0.0004 (950000), DM C vs. L745 Tukey, Figure 2D). L745 application had limited effects on the B-wave amplitude in both conditions, except for the highest intensity flash where the B-wave was significantly elevated (Mixed-effects analysis, p = 0.0325, non-DM C vs. L745 Tukey; 2-way ANOVA p = 0.0271, DM C vs. DM L745 Tukey, Figure 2E), but there were no changes in implicit time to B-wave. On the contrary, we see that rise time in DM conditions is significantly quicker (Mixed-effect analysis, p = 0.0324 (9500); <0.0001 (95000); 0.0001 (950000) DM C vs. DM L745 Tukey; Figure 2G) at every intensity and at the highest intensity for non-DM conditions (Mixed-effects analysis, p = 0.0015, non-DM C vs L745 Tukey; Figure 2G). The faster rise time kinetics in DM retinas suggests that D4Rs contribute to regulating early ON-pathway responses during LA. IN DM, L745 further accelerates rise time to near dark-adapted levels, indicating a reduced capacity to adjust to background illumination and possible machinery issues in dopamine receptors. This suggests that L745 prevents normal LA, supporting a role for D4Rs in mediating this adaptive process.

### 3.3 B-wave amplitudes are elevated in DM retinas after application of D1R antagonist

D1 receptors are localized to the inner nuclear layer (INL), and we wanted to explore ERG changes in non-DM and DM conditions when D1Rs were blocked with SCH-23390 (SCH). The application of SCH had little to no effect on A-wave amplitudes or implicit timing in DA conditions. The B-wave amplitude in DM conditions significantly increased (Mixed-effects analysis, p = 0.0001 (950); 0.0033 (9500), DM C vs. SCH Tukey; Figure 3E) after application of SCH at lower light intensities, but this had no significant effect on implicit timing to the B-wave (Figure 3F). Rise time, however, was significantly delayed in DM conditions after application of SCH (Mixed-effects analysis, p = 0.0003, DM C vs SCH Tukey; Figure 3G). This suggests that DM conditions could have altered receptor properties leading to an increase in overall amplitude.

**Figure 3.**
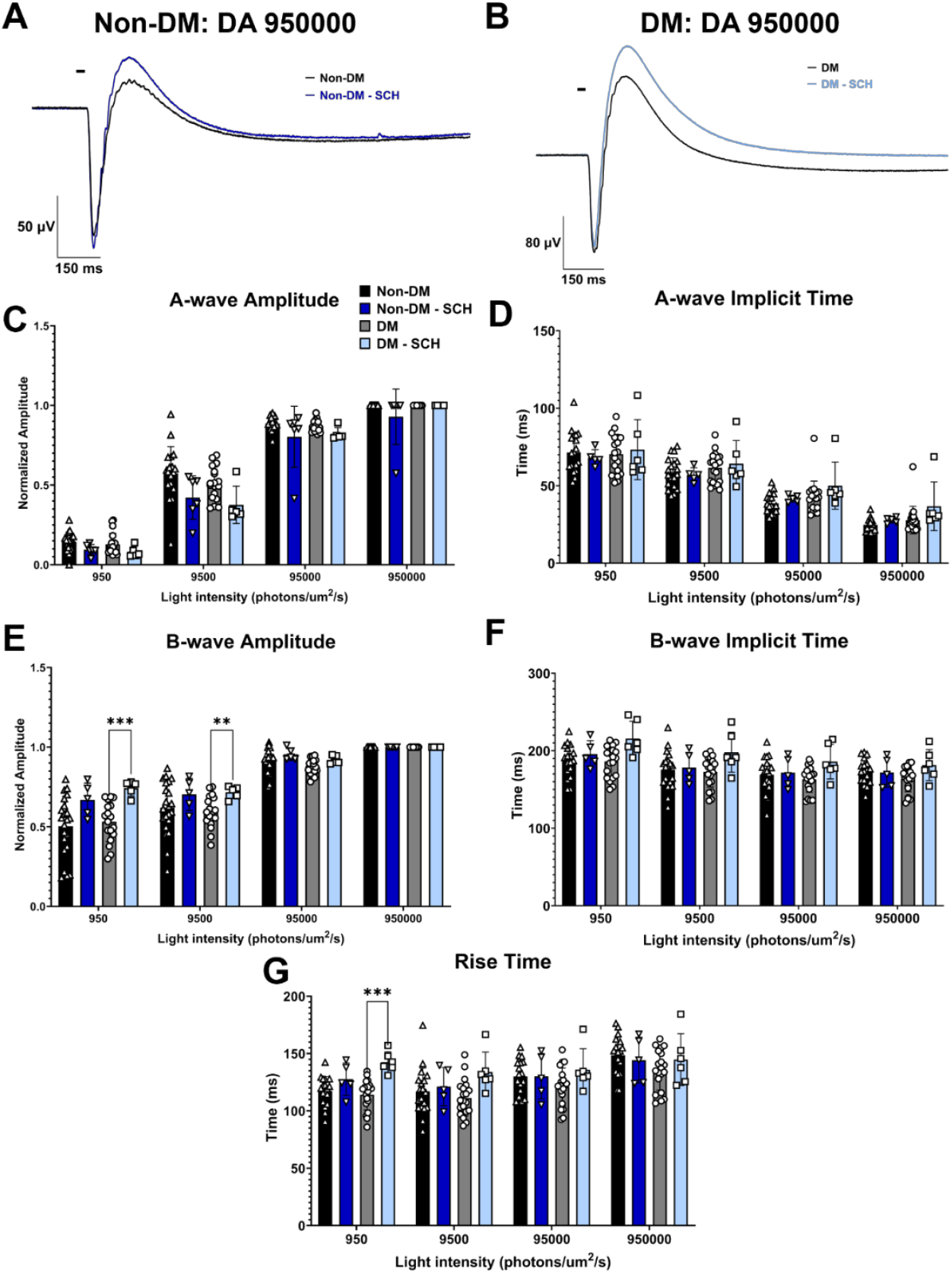
D1 receptor blockade has minimal dark-adapted effects except for increased B-wave amplitude and delayed rise time in diabetic retinas.(A-B) Representative dark-adapted ERG traces from Non-DM and DM retinas before (black) and after SCH-23390 (SCH) application (50 μM; blue and light blue). (C-D) A-wave amplitude and implicit time showed no significant changes after SCH in either group. (D) B-wave amplitude significantly increased in DM retinas at lower flash intensities following SCH application, with no change in Non-DM retinas. (F) B-wave implicit time was unchanged by SCH. (G) Rise time was significantly delayed in DM retinas after D1R blockade, indicating potential deficiencies in inner retinal signaling. Data shown as individual retinas with mean ± SEM; mixed-effects analysis with Tukey corrections.

### 3.4 D1r antagonist has little effect on light adapted ERG responses in DM retinas

As we have done previously, we next wanted to look at how application of SCH affected light adaptation. Neither the A-or B-wave amplitudes had significant changes in either non-DM or DM groups. We saw the biggest change in A-wave implicit timing where the DM group was significantly delayed (Mixed-effects analysis, p = 0.0025, DM C vs SCH Tukey; Figure 4D) which could be due to a slight increase in A-wave amplitude in the DM-SCH group. The rise time got significantly quicker in DM groups after application of SCH at the 95000 photons/μm^2^/s intensity flash (Mixed-effects analysis, p = 0.0114, DM C vs SCH; Figure 4G). This suggests that blockage of D1Rs has a small effect on the ERG’s ability to adjust to increasing background light between groups.

**Figure 4.**
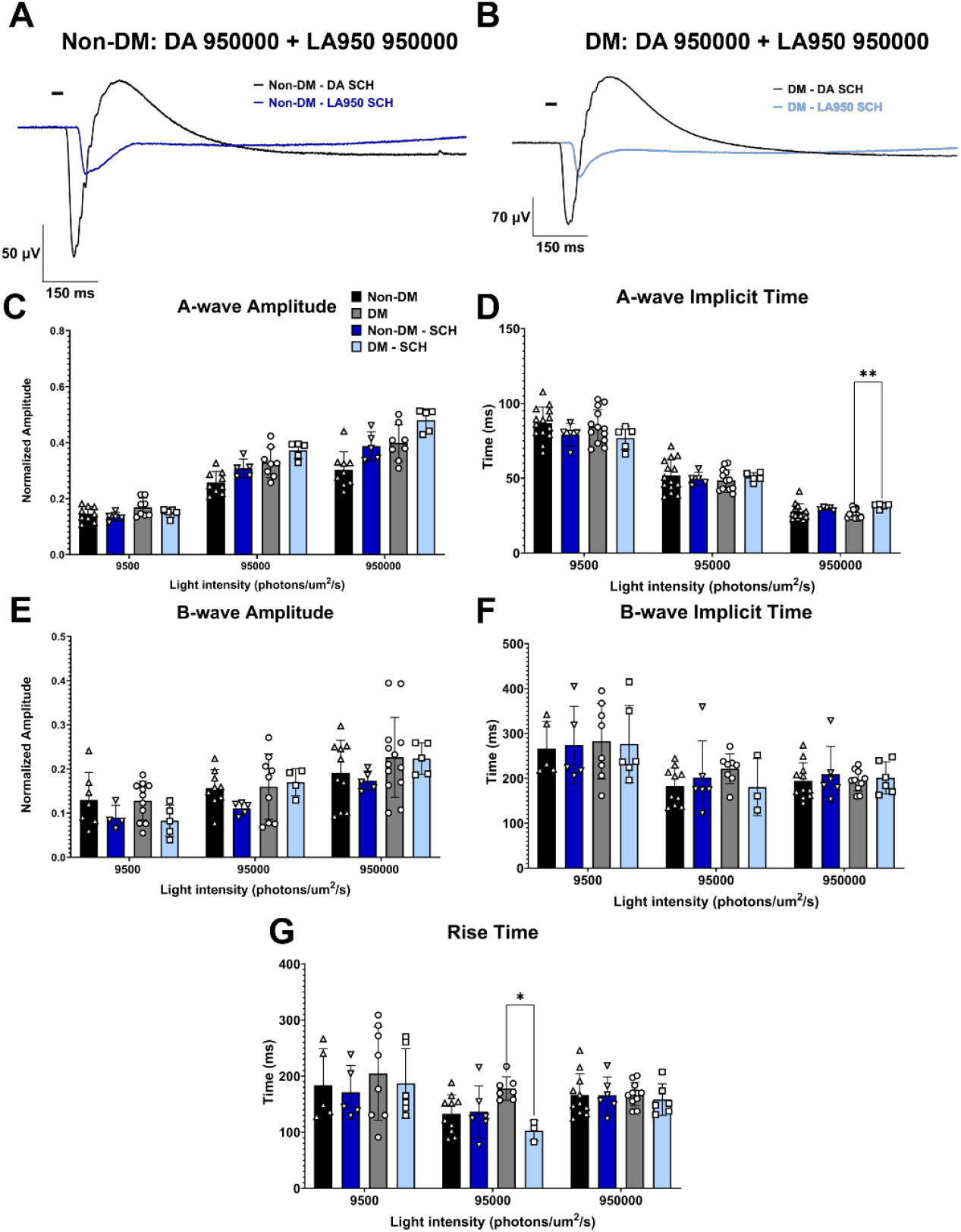
D1 receptor blockade modestly alters light-adaptation timing in diabetic retinas but does not affect ERG amplitudes.(A-B) Representative light-adapted ERG traces from Non-DM and DM retinas before (black) and after SCH application (50 μM; non-DM, blue; DM, light blue). (C, E) A-wave amplitude and B-wave amplitude were unchanged by SCH in both groups. (D) A-wave implicit time was significantly delayed in DM retinas after SCH treatment, consistent with slowed photoreceptor-driven adaptation. (E) B-wave implicit time did not differ across conditions. (G) Rise time was significantly faster in DM retinas at the 95,000 photons/μm^2^/s intensity following SCH application, with no significant changes in Non-DM retinas. Results represent mean ± SEM with individual data points; mixed-effects analysis with Tukey post hoc comparisons.

## 4. DISCUSSION

### 4.1 D4R and D1R antagonist effects on dark-adapted responses in 6-week diabetic retina

ERG changes are some of the first signs of visual deficits perceptible to diabetic complications. Light adaptation is a visual deficit associated with diabetes and dopamine is critical in modulating this process. Therefore, we wanted to investigate what ERG changes would occur after blocking either D1Rs or D4Rs in the retina. In a previous paper, we investigated ERG changes after application of a D4R or D1R agonist in DM and non-DM conditions. We found DM retinas to be non-responsive to either a D1R or D4R agonist and considered DM retinas to have disruptions in dopamine receptor activity or protein levels^[13]^ given there were no mRNA expression changes of either D4Rs or D1Rs. Therefore, we hypothesized that there would be no response change to application of D4R or D1R antagonists in the dark-adapted retina, given our D4R and D1R agonist results. We first investigated what effect L745 (D4R antagonist) would have on dark adapted retinas and found that the A-wave amplitude in diabetic retinas was significantly reduced compared to DM C (Figure 1C). D4Rs are primarily located in the ONL on photoreceptors (Cite) therefore this would explain the reduction in amplitude. If receptor activity is reduced, this could explain the differences observed between non-DM and DM retinas.

Following a similar protocol to L745 experiments, we then looked at the D1R antagonist, SCH. In the dark, we found there to be a significant elevation in b-wave amplitude in DM compared to non-DM conditions (p = 0.0001, Figure 3E). D1Rs are located in the INL, therefore we would expect to see changes here (if any).

### 4.2 D4R and D1R antagonist effects on light adaptation in the 6-week diabetic retina

Light adaptation is a critical feature of the retina and dopamine is the regulator of that process. We investigated how blocking dopamine receptors would affect the light adapted process. Following the same flow as dark-adapted recordings, we looked at blocking D4Rs with L745. We found the light-adapted A-wave amplitude in both the non-DM and DM conditions to be significantly elevated (Figure 2C). We attributed the blockage of D4Rs as disabling the ability for the retina to adequately adapt to increasing background light. When there is enough light that goes beyond the adequate level for rods to respond, there must be a mechanism to saturate the rods so the rod pathway is less active. The b-wave amplitude is also significantly elevated with application of L745 at the highest intensity flash (Figure 2E) in DM, therefore this points to a cellular machinery issue similar to ON-s ganglion cells inefficient signaling with the application of a D4R agonist^[31]^. A more drastic change can be seen with timing where both non-DM and DM conditions were significantly delayed (Figure 2D) with application of L745, even more-so in the DM condition at the lower light intensity. Lastly, the rise time is significantly reduced (Figure 2G) pointing to a quicker transition from a-wave to b-wave. Blocking D1Rs with SCH did not show any significant changes on light adapted ERG signaling besides a slight elevation in A-wave implicit time (Figure 4D) and rise time (Figure 4G). The critical part of light adaptation present in the ERGs likely depends on D4Rs; when these are dysfunctional, as seen in D4R knockout mice, the retina shows a reduced ability to adapt^[32]^. Reductions in D4R activity disrupt cAMP regulation in photoreceptors^[32]^, which is likely contributing to consistently delayed kinetics in light adaptation. Therefore, it is plausible that there is more adenylyl cyclase activity (AC1), causing more cAMP accumulation, thus increasing Ca^2+^ influx.

## 6. Conclusions

ERG recordings reveal subtle deficits in non-DM vs DM conditions, but there are clear deficiencies in dopamine signaling in diabetes as early as 6-weeks in rodents. Blocking D4Rs reveals diabetic-dependent changes in ERG signaling, particularly in A-wave amplitude, timing, and light adaptation ability. D1R produces subtle but significant changes in B-wave amplitude in the dark, but no statistically relevant change in light adaptation, suggesting D4Rs are potentially altered. Together, these results highlight D4R signaling as a potential site of vulnerability in early diabetes as it relates to light adaptation and dopamine signaling. Thus, targeting D4R signaling pathways may provide therapeutic potential to preserve light adaptation.

## References

1. Lundeen EA. B.-C.Z., Rein DB, Wittenborn JS, Saaddine J, Lee AY, Flaxman AD., Prevalence of Diabetic Retinopathy in the US in 2021. JAMA Ophthalmology, 2023. 141(8): p. 747–754.

2. Shyam, M., et al., Diabetic retinopathy: a comprehensive review of pathophysiology and emerging treatments. Molecular Biology Reports, 2025. 52(1).

3. Eggers, E.D., Visual Dysfunction in Diabetes. Annual Review of Vision Science, 2023. 9(Volume 9, 2023): p. 91–109.

4. Eggers, E.D. and T.A. Carreon, The effects of early diabetes on inner retinal neurons. Vis Neurosci, 2020. 37: p. E006.

5. Cusick, M., et al., Central Visual Function and the NEI-VFQ-25 Near and Distance Activities Subscale Scores in People with Type 1 and 2 Diabetes. American Journal of Ophthalmology, 2005. 139(6): p. 1042–1050.

6. Klein, R., B.E. Klein, and S.E. Moss, Visual impairment in diabetes. Ophthalmology, 1984. 91(1): p. 1–9.

7. Klein, R., et al., The NEI-VFQ-25 in People With Long-term Type 1 Diabetes Mellitus: The Wisconsin Epidemiologic Study of Diabetic Retinopathy. Archives of Ophthalmology, 2001. 119(5): p. 733–740.

8. Akkaya, S., et al., National Eye Institute Visual Function Scale in Type 2 Diabetes Patients. Journal of Ophthalmology, 2016. 2016(1): p. 1549318.

9. Aung, M.H., et al., Early Visual Deficits in Streptozotocin-Induced Diabetic Long Evans Rats. Investigative Ophthalmology & Visual Science, 2013. 54(2): p. 1370–1377.

10. Layton, C.J., R. Safa, and N.N. Osborne, Oscillatory potentials and the b-Wave: Partial masking and interdependence in dark adaptation and diabetes in the rat. Graefes Archive for Clinical and Experimental Ophthalmology, 2007. 245(9): p. 1335–1345.

11. Han, Y., et al., Multifocal Electroretinogram Delays Predict Sites of Subsequent Diabetic Retinopathy. Investigative Ophthalmology & Visual Science, 2004. 45(3): p. 948–954.

12. Bearse, M.A., Jr., et al., Retinal function in normal and diabetic eyes mapped with the slow flash multifocal electroretinogram. Invest Ophthalmol Vis Sci, 2004. 45(1): p. 296–304.

13. MacIsaac, A.R., et al., Impaired dopamine signaling in early diabetic retina: Insights from D1R and D4R agonist effects on whole retina responses. Experimental Eye Research, 2024. 247: p. 110049.

14. Aung, M.H., et al., Dopamine Deficiency Contributes to Early Visual Dysfunction in a Rodent Model of Type 1 Diabetes. The Journal of Neuroscience, 2014. 34(3): p. 726.

15. Kim, M.K., et al., Dopamine Deficiency Mediates Early Rod-Driven Inner Retinal Dysfunction in Diabetic Mice. Investigative Ophthalmology & Visual Science, 2018. 59(1): p. 572–581.

16. Motz, C.T., et al., Novel Detection and Restorative Levodopa Treatment for Preclinical Diabetic Retinopathy. Diabetes, 2020. 69(7): p. 1518–1527.

17. Witkovsky, P., Dopamine and retinal function. Doc Ophthalmol, 2004. 108(1): p. 17–40.

18. Roy, S. and G.D. Field, Dopaminergic modulation of retinal processing from starlight to sunlight. J Pharmacol Sci, 2019. 140(1): p. 86–93.

19. Jackson, C.R., et al., Retinal dopamine mediates multiple dimensions of light-adapted vision. J Neurosci, 2012. 32(27): p. 9359–68.

20. Farshi, P., et al., Dopamine D1 receptor expression is bipolar cell type-specific in the mouse retina. Journal of Comparative Neurology, 2016. 524(10): p. 2059–2079.

21. Cohen, A.I., et al., Photoreceptors of mouse retinas possess D4 receptors coupled to adenylate cyclase. Proceedings of the National Academy of Sciences, 1992. 89(24): p. 12093–12097.

22. Derouiche, A. and E. Asan, The dopamine D2 receptor subfamily in rat retina: ultrastructural immunogold and in situhybridization studies. European Journal of Neuroscience, 1999. 11(4): p. 1391–1402.

23. Li, H., et al., Adenosine and dopamine receptors coregulate photoreceptor coupling via gap junction phosphorylation in mouse retina. Journal of Neuroscience, 2013. 33(7): p. 3135–3150.

24. Laura, L.K., et al., Localization and regulation of dopamine receptor D4 expression in the adult and developing rat retina. Experimental Eye Research, 2008. 87(5): p. 471–477.

25. Veruki, M.L., Dopaminergic Neurons in the Rat Retina Express Dopamine D2/3 Receptors. European Journal of Neuroscience, 1997. 9(5): p. 1096–1100.

26. Veruki, M.L. and H. Wässle, Immunohistochemical Localization of Dopamine D Receptors in Rat Retina. European Journal of Neuroscience, 1996. 8(11): p. 2286–2297.

27. Tran, V.T. and M. Dickman, Differential localization of dopamine D1 and D2 receptors in rat retina. Investigative Ophthalmology & Visual Science, 1992. 33(5): p. 1620–1626.

28. Tuyen, V., et al., Radioimmunoligand characterization and immunohistochemical localization of dopamine D2 receptors on rods in the rat retina. Brain Research, 1993. 614(1): p. 57–64.

29. Flood, M.D. and E.D. Eggers, Dopamine D1 and D4 receptors contribute to light adaptation in ON-sustained retinal ganglion cells. Journal of Neurophysiology, 2021. 126(6): p. 2039–2052.

30. Flood, M.D., et al., Early diabetes impairs ON sustained ganglion cell light responses and adaptation without cell death or dopamine insensitivity. Exp Eye Res, 2020. 200: p. 108223.

31. Flood, M.D., A.J. Wellington, and E.D. Eggers, Impaired Light Adaptation of ON-Sustained Ganglion Cells in Early Diabetes Is Attributable to Diminished Response to Dopamine D4 Receptor Activation. Investigative Ophthalmology & Visual Science, 2022. 63(1): p. 33–33.

32. Nir, I., et al., Dysfunctional Light-Evoked Regulation of cAMP in Photoreceptors and Abnormal Retinal Adaptation in Mice Lacking Dopamine D4 Receptors. The Journal of Neuroscience, 2002. 22(6): p. 2063–2073.

